# Reallocation to lateral and early-emerging axial roots allows maize (*Zea mays L.*) with reduced nodal root number to more efficiently forage for nitrate

**DOI:** 10.1101/533042

**Authors:** Haichao Guo, Larry M. York

## Abstract

Previous simulations indicated reduced nodal root number (NRN) was promising for maize (*Zea mays* L.) breeding, and were partially confirmed using variation in NRN among inbreds. However, the exact mechanism was unknown, therefore manipulative experiments were conducted in hydroponics and tall solid-media mesocosms with treatments involving no nodal root excision (0% NRE) or excising either 33% or 67% of the nodal roots (NR) as they emerged under high or low levels of nitrogen (N). Reduced NRN was hypothesized to increase elongation of all remaining root classes, increase N acquisition under low N, and increase shoot mass. In both experiments, plants with 67% NRE had 12% and 19% less root fraction of total biomass, 61% and 91% greater lateral-to-axial root length ratio regardless of N levels; and 61% and 182% greater biomass of embryonic roots under low N, compared to 0% NRE for hydroponics and mesocosms studies, respectively. In hydroponics, regardless of NRE level, specific root respiration under high N was 2.6 times of low N, and was greatest at depth. Under low N in mesocosms, plants with 67% NRE had 52% greater shoot biomass, 450% greater root length at depth, and 232% greater deep-injected ^15^N content in the shoot relative to 0% NRE, however biomass in hydroponics did not differ based on NRE. These results reveal the mechanism by which plants with fewer nodal roots increase N capture and shoot mass by reallocation of biomass to lateral, embryonic, and first whorl nodal roots that increases foraging efficiency in solid media.

**Summary:** Reallocating root biomass from nodal roots to lateral and early-emerging axial roots allows grasses to capture more nitrogen under limiting conditions, including by increasing foraging at depth.

## INTRODUCTION

Roots are the major interface between plants and the soil, with a key function to extract nutrients and water that are required for plant productivity (Smith and De Smet, 2012; Meister et al., 2014). Plants have evolved the ability to proliferate roots in response to heterogeneity of resources, belowground competition, and the inevitability of root loss (Lynch, 2018). Plants respond to resource availability by adjusting their relative shoot and root mass allocation to maximize their relative growth rate (Ågren and Franklin, 2003), and generally allocate relatively more biomass to roots if the limiting factor for growth is a soil resource (Poorter et al., 2012). The fraction of newly fixed carbon from photosynthesis allocated to roots can exceed 50%, and the proportion to roots significantly increases under edaphic stress (Lambers et al., 1996; Rachmilevitch et al., 2015). Root phenes are the general elemental units of phenotype (Lynch and Brown, 2012) influencing resource acquisition, while phene states represent the particular value a phene has taken (York et al., 2013). Studies have shown that root phene states that reduce the metabolic cost of soil exploration permit greater root growth, which improves the capture of deep nitrate (Chimungu et al., 2014; Saengwilai et al., 2014), deep water (Gao and Lynch, 2016), and shallow phosphorus (Strock et al., 2018). ‘Cheap root’ phene states are those that reduce the metabolic burden of the root system with benefits for plant performance, including decreased root diameter, increased allocation to cheaper root classes such as lateral roots and root hairs, reduced root cortical cell area, and reduction in root secondary growth in dicots (Lynch, 2013; Meister et al., 2014; Galindo-Castaneda et al., 2018; Strock et al., 2018).

Maize plays an important role in the global food supply and industrial production, including starch, sweeteners, oil, beverages, glue, industrial alcohol, and fuel ethanol (Ranum et al., 2014). Breeding for optimized maize root system architecture is required to obtain maximum grain yield while reducing nutrient leaching and improving drought resistance in high-input agroecosystems by increasing soil resource acquisition efficiency (Mi et al., 2016; Lynch, 2018). The mature maize root system consists of the embryonic root (ER) and postembryonic root systems. The embryonic primary and seminal roots together with their laterals are important for maize seedling vigor during early development, but the postembryonic shoot-borne nodal roots (NR) and their laterals become the dominant root system by mass as the crop matures (Hochholdinger, 2009; Hochholdinger et al., 2018). Nodal roots that are formed belowground are called crown roots, whereas those that are formed aboveground are designated brace roots (Hochholdinger, 2009; Saengwilai et al., 2014). Evolution of root system architecture towards more nitrogen-efficient states corresponds to historical maize yield trends and increased tolerance of nitrogen (N) stress in the U.S. (Hammer et al., 2009; York et al., 2015).

The ‘steep, deep, cheap’ (SCD) ideotype of the maize root system has been proposed for optimal N and water foraging, and consists of root architectural, anatomical, and physiological phene states to increase rooting depth and N and water use efficiency in specific environments (Lynch, 2013; York and Lynch, 2015). Nodal root (NR) number is an aggregate phene consisting of the number of nodal whorls and the number of roots per whorl (Saengwilai et al., 2014; York and Lynch, 2015), and flowering time played a key role in shaping NR number via indirect selection during maize domestication (Zhang et al., 2018). The functional-structural plant model *SimRoot* predicted that reduced NR number could enhance maize growth under low levels of N by increasing growth of lateral roots and N acquisition (York et al., 2013; York, 2014). Field and greenhouse studies later confirmed maize genotypes with fewer nodal roots had greater rooting depth and ability to acquire N from deep soil of low N soils (Saengwilai et al., 2014), and improved water acquisition from drying soil (Gao and Lynch, 2016). However, these studies have relied on intraspecific diversity to create root phene variation, which could be confounded by the multitude of differences that potentially exist among genotypes that were not necessarily measured or reported (Lynch, 2011). In this study, we used root excision to manipulate NR number of maize inbred line B73 grown in growth chamber hydroponics and in greenhouse tall mesocosms to explore the effect of reduced NR number on N acquisition. The objective of this study was using maize plants with the same genetic and phenotypic background to test the hypotheses that reducing NR number would (1) increase elongation of embryonic roots (ER) and the remaining NR and their laterals due to reallocation of carbon, (2) increase shoot biomass in all conditions due to reallocating carbon to the shoot, and (3) increase N acquisition in low N environments.

## RESULTS

### Growth chamber hydroponics experiment

For whole-plant phenes in the hydroponics experiment, the main effects of N treatment and nodal root excision (NRE) treatment were all significant for NR number, assimilation rate, specific root length, and axial root length. Shoot biomass, root biomass, total biomass, transpiration rate, stomatal conductance, total root length, specific root respiration (SRR) by length, SRR by mass and lateral root length were only affected by N treatment, and root mass fraction and lateral-to-axial root length ratio were only affected by NRE treatment (Supplemental Table S1). Low N supply substantially reduced NR number by 13% (Fig. 1A), shoot biomass by 38% (Fig. 1B), root biomass by 30% (Fig. 1C), total biomass by 37% (Fig. 1D), assimilation rate by 59% (Fig. 1F), transpiration rate by 71% (Fig. 1G), stomatal conductance by 79% (Fig. 1H), total root length by 46% (Fig. 1I), specific root length by 25% (Fig. 1J), specific root respiration by root length by 50% (Fig. 1K), specific root respiration by mass by 62% (Fig. 1L), axial root length by 35% (Fig. 1M), and lateral root length by 46% (Fig. 1N) at 24 days after planting (DAP). 67% NRE led to 52% reduction in NR number (Fig. 1A), 12% reduction in root mass fraction (Fig. 1E), 10% reduction in assimilation rate (Fig. 1F), 29% reduction in axial root length (Fig. 1M), and 61% increase in lateral-to-axial root length ratio (Fig. 1O). NR excision had no effect on NR emergence time (Data not shown). No significant interactions between N treatment and NR excision treatment were observed for the phenes measured (Supplemental Table S1).

**Figure 1.**
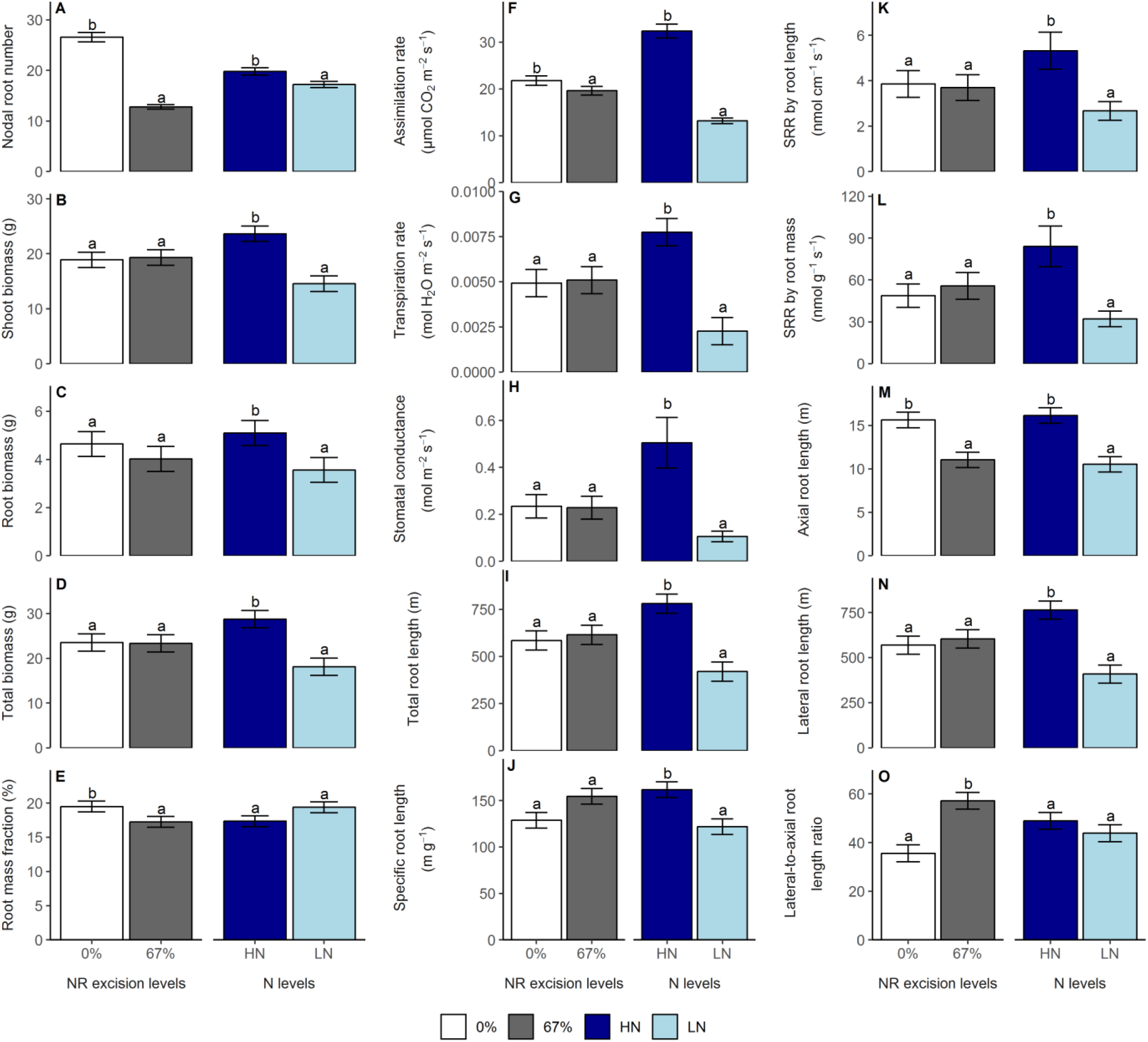
Main effects of N treatment and NR excision treatment on whole-plant shoot and root phenes in the growth chamber hydroponics study. Each column represents the mean of n=5 for each treatment. Error bars represent standard errors, and columns with the same letter under same main effect were not significantly different at p≤0.05 according to Tukey’s test. SRR: Specific root respiration.

Specific root classes are referred to as ER and NR, which are further grouped by whorl with appended numbers referring to order of emergence; therefore NR1 is the first whorl of NR that emerges. Under high N conditions, 67% NRE reduced root biomass by 74% (Fig. 2A), root respiration by 74% (Supplemental Fig. S1A), root length by 68% (Supplemental Fig. S1B), and axial root length by 72% (Supplemental Fig. S1F) in NR1; and led to 128% increase of specific root length (Supplemental Fig. S1C) and 55% reduction of SRR by length in NR5 (Supplemental Fig. S1D). A trend of greater root biomass in ER induced by 67% NRE was observed under high N conditions. Under low N conditions, 67% NRE increased root biomass by 61% (Fig. 2B) and root respiration by 100% (Supplemental Fig. S1I) in ER, reduced root biomass by 47% (Fig. 2B) and axial root length by 69% (Supplemental Fig. S1N) in NR1, led to 67% reduction of axial root length (Supplemental Fig. S1N) and 156% increase of lateral-to-axial root length ratio in NR2 (Supplemental Fig. S1P), increased specific root length by 52% (Supplemental Fig. S1K) and lateral-to-axial root length ratio by 169% (Supplemental Fig. S1P) in NR3, and increased specific root length by 79% (Supplemental Fig. S1K), lateral root length by 222% (Supplemental Fig. S1O) and lateral-to-axial root length ratio by 267% (Supplemental Fig. S1P) in NR4. 67% NRE had no impact on SRR by root mass across all root classes under both N levels (Supplemental Fig. S1E and 1M).

**Figure 2.**
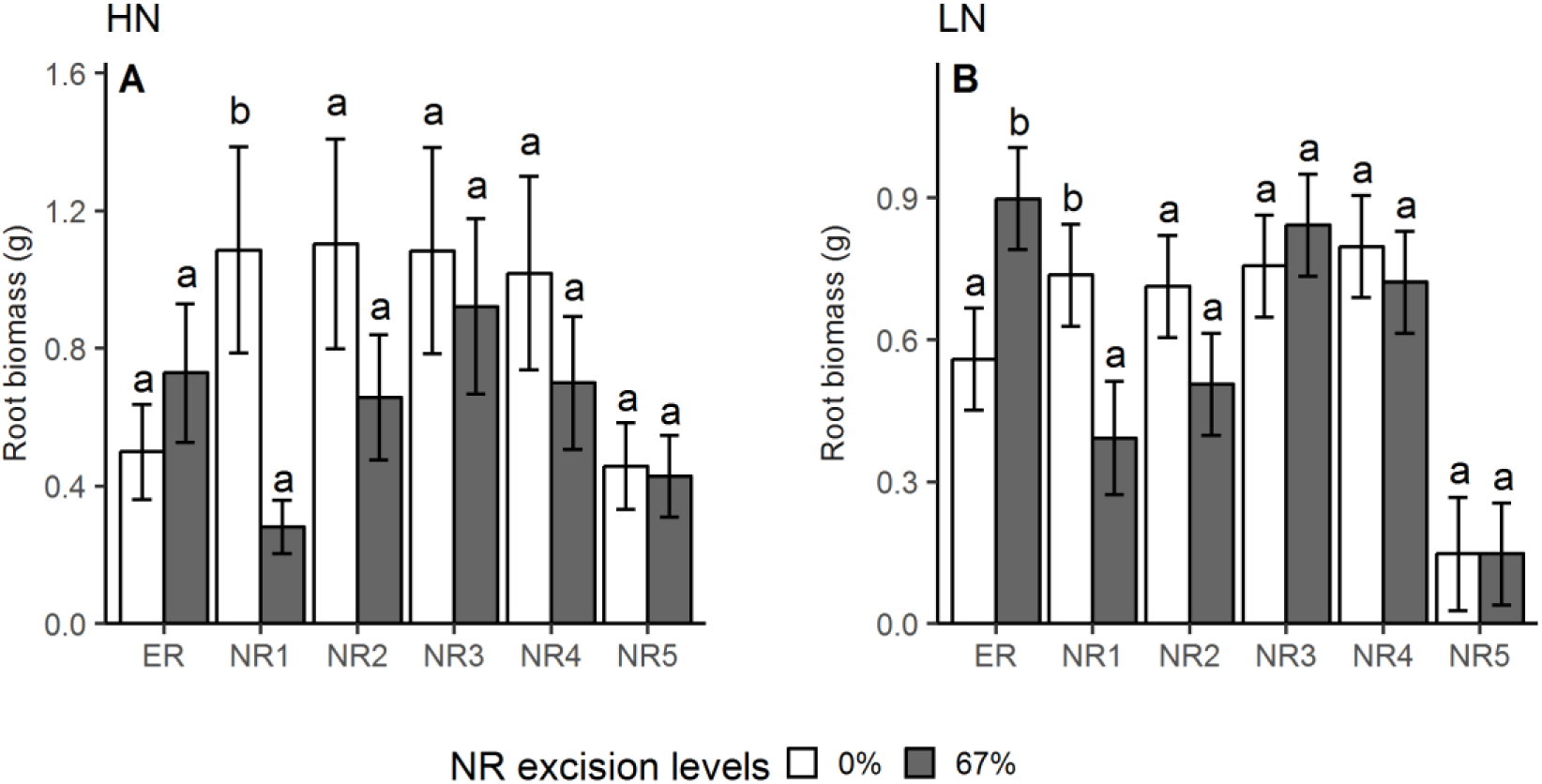
Effect of NR excision treatment on root biomass of different root classes under high N (A) and low N (B) conditions in growth chamber hydroponics study. Each column represents the mean of n=5 for each treatment. Error bars represent standard errors, and columns with the same letter under same root class were not significantly different at p≤0.05 according to Tukey’s test. ER: embryonic roots, NR1: 1^st^ whorl nodal roots, NR2: 2^nd^ whorl nodal roots, NR3: 3^rd^ whorl nodal roots, NR4: 4^th^ whorl nodal roots, NR5: 5^th^ whorl nodal roots.

Under high N conditions, 67% NRE led to 55% reduction of root biomass (Supplemental Fig. S2A) and 47% increase of SRR by mass at the depth of 90-120 cm (Supplemental Fig. S2F). Under low N conditions, 67% NRE reduced root respiration by 40% in 60-90 cm and by 43% in 90-120 cm (Supplemental Fig. S2K), reduced SRR by length by 43% in 60-90 cm, reduced axial root length by 33% in 0-30 cm, by 51% in 30-60 cm, by 62% in 60-90 cm and by 53% in 90-120 cm (Supplemental Fig. S2P), and led to 48% increase of specific root length and 196% increase of lateral-to-axial root length ratio in 60-90 cm (Supplemental Fig. S2R). NR excision had no effect on root length and lateral root length across all depths under both N levels (Supplemental Fig. S2C, 2L, 2H and 2Q). Regardless of NR excision levels, SRR by mass increased with at the greatest depth under both N levels (Fig. 3).

**Figure 3.**
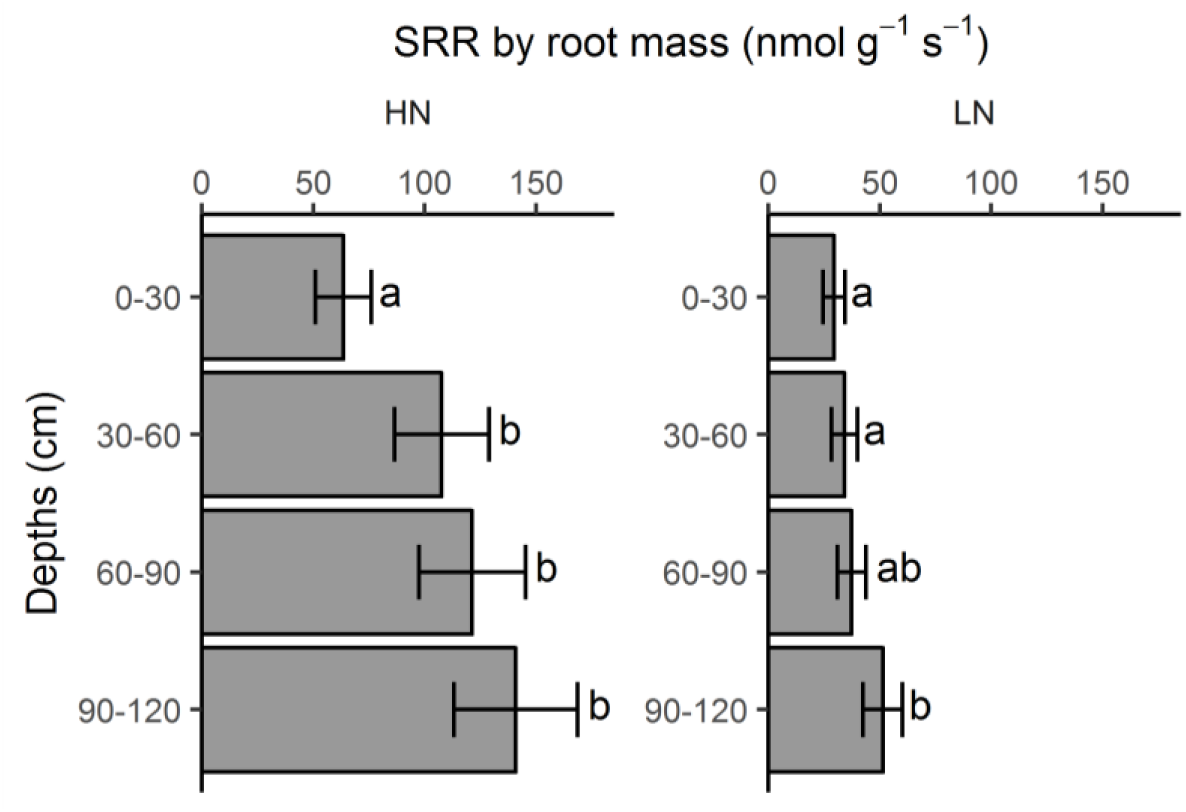
Effects of rooting depths on SRR by root mass under high N and low N conditions in growth chamber hydroponics study. Each column represents the mean of n=10 for each depth. Error bars represent standard errors, and columns with the same letter under same N level were not significantly different at p≤0.05 according to Tukey’s test. SRR: Specific root respiration.

### Greenhouse mesocosms experiment

For whole-plant shoot and root phenes in mesocosms experiment, the main effects of N treatment and NR excision treatment were all significant for NR number, shoot biomass, root mass fraction, specific root length, and lateral-to-axial root length ratio. Root biomass, total biomass, shoot N concentration, shoot ^15^N concentration, total root length, and lateral root length were only affected by N treatment (Supplemental Table S1). Low N supply reduced NR number by 22% (Fig. 4A), shoot biomass by 68% (Fig. 4B), root biomass by 59% (Fig. 4C), total biomass by 66% (Fig. 4D), shoot N concentration by 43% (Supplemental Fig. S3A), total root length by 48% (Fig. 4H), lateral root length by 50% (Fig. 4J) and lateral-to-axial root length by 29% (Fig. 4L), and increased root mass fraction by 23% (Fig. 4E), shoot ^15^N concentration by 101% (Supplemental Fig. S3B) and specific root length by 21% (Fig. 4K) at 42 DAP. The 33% NRE level decreased NR number by 34% (Fig. 4A), increased specific root length by 25% (Fig. 4K) and lateral-to-axial root length ratio by 65% (Fig. 4L), and had no impact on shoot biomass (Fig. 4B) and root mass fraction (Fig. 4E) compared to 0% NRE across all N levels. Increasing NRE level to 67% decreased NR number by 60% (Fig. 4A) and root mass fraction by 19% (Fig. 4E), and increased shoot biomass by 35% (Fig. 4B), specific root length by 41% (Fig. 4K) and lateral-to-axial root length ratio by 91% (Fig. 4L), as compared with 0% NRE across all N levels. 67% NRE led to 5^th^ whorl of NR emerging 2 days earlier and 6^th^ whorl of NR emerging 8 days earlier compared to 0% NRE under high N level, respectively (Fig. 5). The interactions of N treatment and NRE treatment had significant effects on ^15^N content in shoot, N content in shoot, and axial root length (Supplemental Table S1). The 33% and 67% NRE levels significantly increased N content in shoot by 48% and 53%, and ^15^N content in shoot by 162% and 232% under low N, respectively, but these was not observed under high N (Fig. 4F and 4G). Also, both excision levels significantly decreased axial root length under high N, but this was not observed under low N (Fig. 4I).

**Figure 4.**
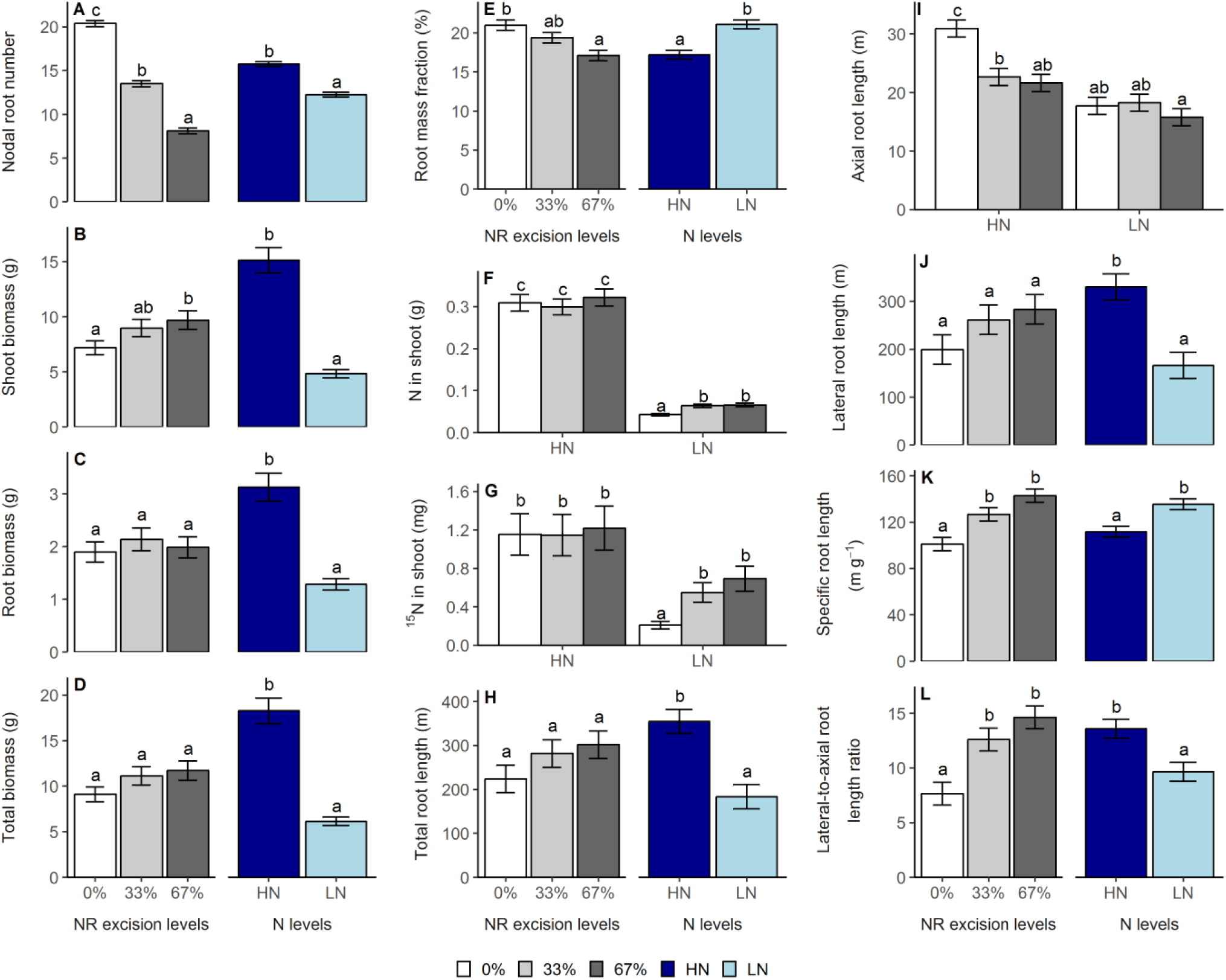
Main effects of N treatment and NR excision treatment (A, B, C, D, E, H, J, K, and L) and their interaction effects (F, G and I) on whole-plant shoot and root phenes in the greenhouse mesocosms study. Each column represents the mean of n=4 for each treatment. Error bars represent standard errors, and columns with the same letter under same main effect were not significantly different at p≤0.05 according to Tukey’s test.

**Figure 5.**
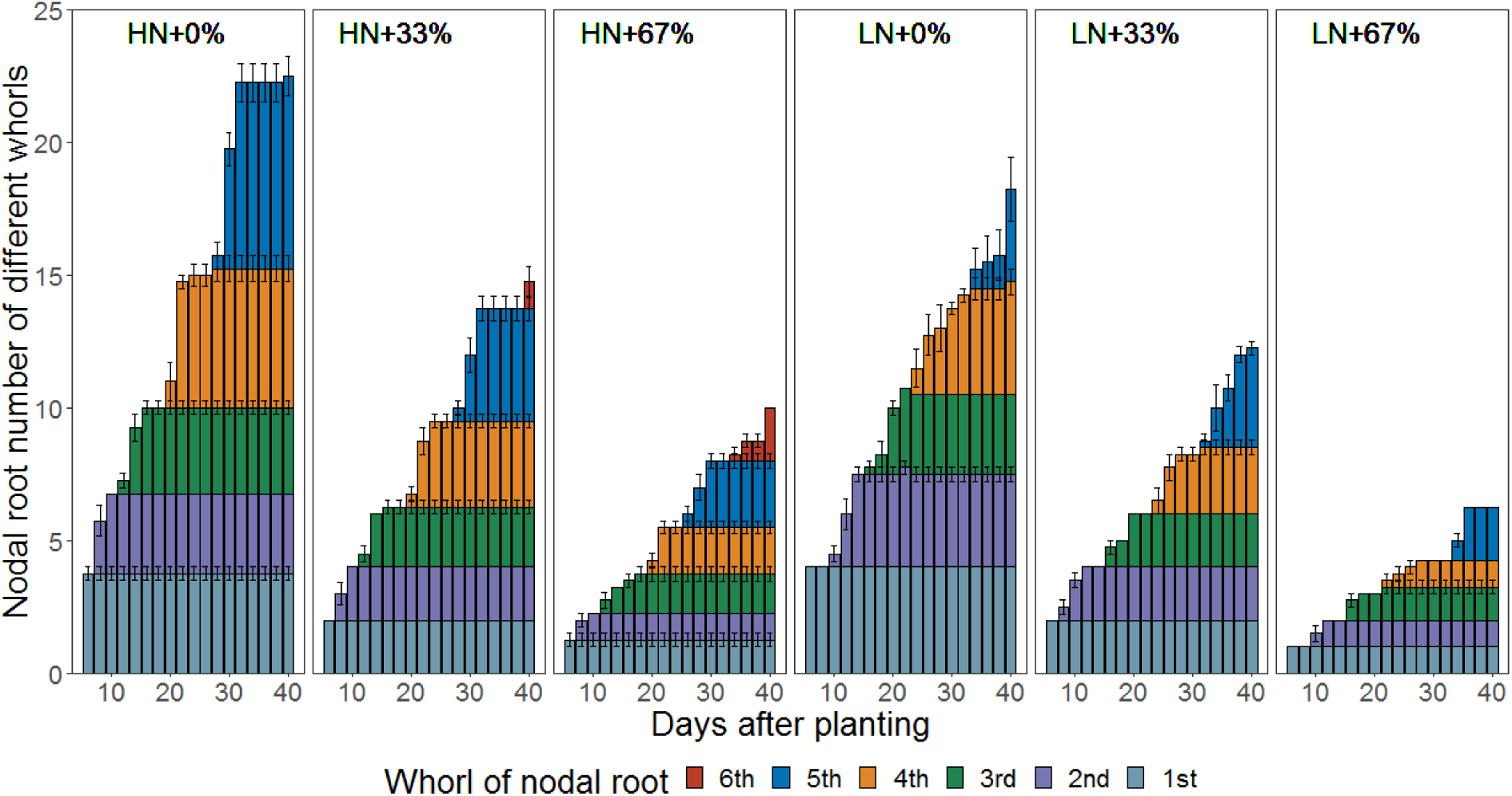
NR number of different whorls with time in the greenhouse mesocosms. Each column with the same color represents the mean of n=4 for each treatment. Error bars represent standard errors. HN+0%: 0% NR excision under high N. HN+33%: 33% NR excision under high N. HN+67%: 67% NR excision under high N. LN+0%: 0% NR excision under low N. LN+33%: 33% NR excision under low N. LN+67%: 67% NR excision under low N.

Under high N conditions, 67% NRE increased root biomass by 112% and individual axial root length by 94% in ER, decreased root biomass by 33% in NR4 and by 34% in NR5 (Fig. 6A), led to 109% increase of axial root length in ER, decreased axial root length by 60% in NR1, 67% in NR2, 63% in NR3, 58% in NR4 and 53% in NR5 (Supplemental Fig. 4B), and increased lateral-to-axial root length ratio by 245% in NR2 and 260% in NR5 (Supplemental Fig. 4E). Under low N conditions, 67% NRE increased root biomass by 182% in ER (Fig. 6B), increased individual axial root length by 164% in ER and by 148% in NR3 (Fig. 6D), and increased lateral-to-axial root length ratio by 279% in NR4 (Supplemental Fig. 4J). Removal of NR had no impact on root length and lateral root length across all root classes under both N levels (Supplemental Fig. S4A, S4F, S4C and S4H).

**Figure 6.**
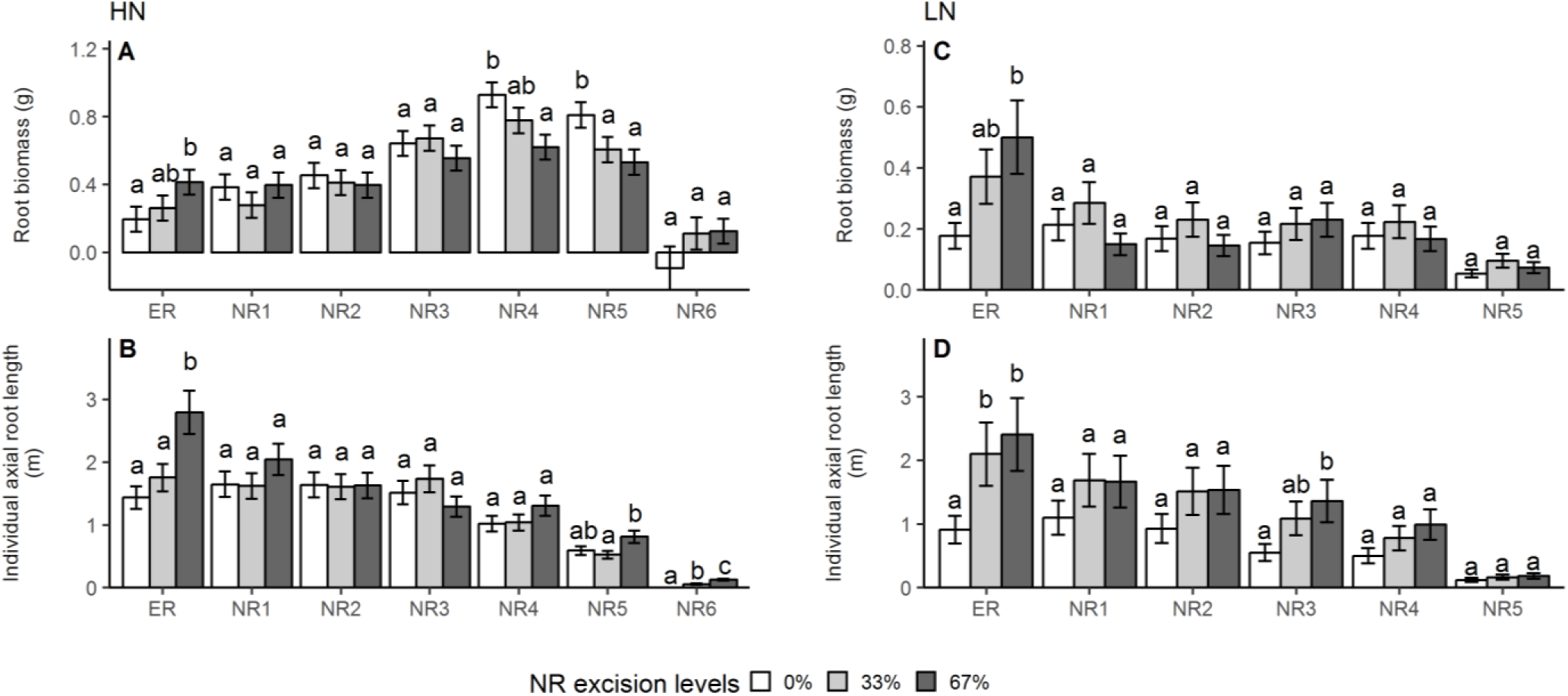
Effect of NR excision treatment on root biomass (A, B) and individual axial root length (C, D) of different root classes under high N and low N conditions in the greenhouse mesocosms. Each column represents the mean of n=4 for each treatment. Error bars represent standard errors, and columns with the same letter under same root class were not significantly different at p≤0.05 according to Tukey’s test. ER: embryonic roots, NR1: 1^st^ whorl nodal roots, NR2: 2^nd^ whorl nodal roots, NR3: 3^rd^ whorl nodal roots, NR4: 4^th^ whorl nodal roots, NR5: 5^th^ whorl nodal roots, NR6: 6^th^ whorl nodal roots.

Under high N conditions, 67% NRE increased root length by 59% and lateral root length by 71% in 120-150 cm (Fig. 7A), decreased axial root length by 48% in 0-30 cm by 42% in 30-60 cm, by 31% in 60-90 cm, by 36% in 90-120 cm (Supplemental Fig. S5B), increased specific root length by 48% in 0-30 cm by 31% in 30-60 cm, by 33% in 120-150 cm (Supplemental Fig. S5D), and increased lateral-to-axial root length by 133% in 0-30 cm by 123% in 30-60 cm, by 82% in 90-120 cm (Supplemental Fig. S5E). Under low N conditions, 67% NRE increased root length by 450% (Fig. 7B), lateral root length by 548% and root biomass by 228% in 120-150 cm (Supplemental Fig. S5H and 5F), led to decrease of axial root length by 52% in 0-30 cm, by 46% in 30-60 cm and 130% increase in 120-150 cm (Supplemental Fig. S5G), increased specific root length by 53% in 0-30 cm, by 67% in 120-150 cm (Supplemental Fig. S5I), and increased lateral-to-axial root length by 205% in 0-30 cm by 114% in 30-60 cm, by 143% in 60-90 cm by 178% in 120-150 cm (Supplemental Fig. S5J).

**Figure 7.**
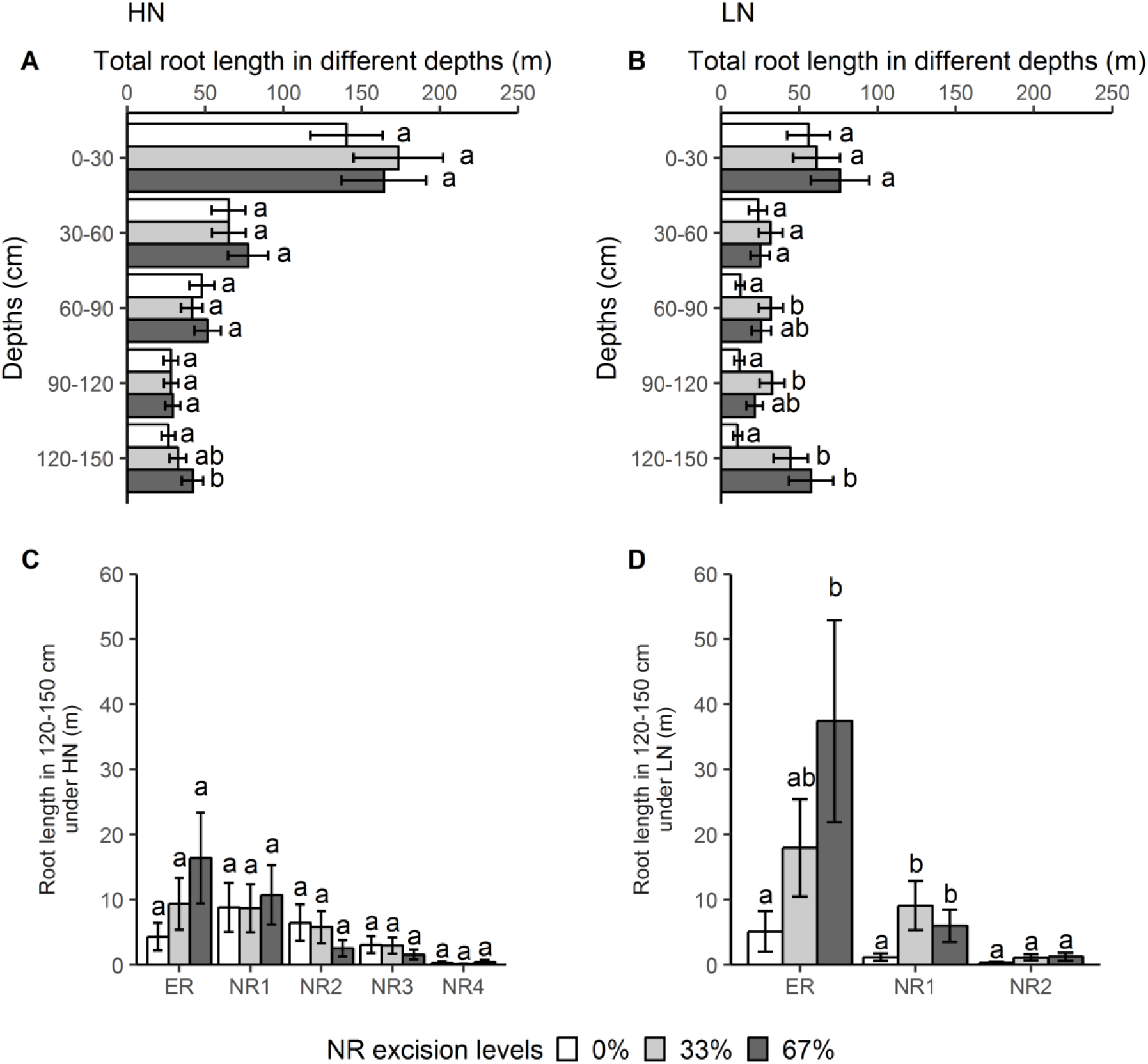
Effect of NR excision treatment on root length distribution across rooting depths under high N level (A) and low N (B) conditions, and root length of different root classes in 120-150 cm (C, D) in the greenhouse mesocosms. Each column represents the mean of n=4 for each treatment. Error bars represent standard errors, and columns with the same letter under same depth or root class were not significantly different at p≤0.05 according to Tukey’s test.

ER and the first four whorls of nodal roots under high N conditions, and ER and the first two whorls of nodal roots under low N conditions reached the depth of 120-150 cm at 42 DAT. In depth of 120-150 cm, 67% NRE increased root length by 630% in ER and by 690% in NR1 under low N conditions (Fig. 7D), and a trend of greater root length in ER induced by 67% NRE was also observed under high N conditions (Fig. 7C).

## DISCUSSION

Reducing the construction and maintenance metabolic costs of roots has been proposed as a strategy for crop breeding to increase N and water use efficiency (Lynch, 2013, 2018). In this study, root system architecture (RSA) was manipulated in order to reduce the number of nodal roots (NR) to test hypotheses that this would influence RSA, increase shoot biomass in all conditions, and increase nitrogen (N) uptake in low N conditions. In hydroponics and mesocosms, 67% nodal root excision (NRE) led to increased embryonic root mass accumulation and greater overall lateral-to-axial root ratio, indicating an inherent reallocation of root system carbon to other root sinks. However, contrary to our hypothesis, 67% NRE had no effect on shoot biomass or shoot N content in either hydroponics N level, but increased shoot biomass by 35% across all N levels in mesocosms. Remarkably, under low N in mesocosms, 67% NRE allowed the proliferation of 450% more root length at the deepest depth, increased deep-injected ^15^N content in the shoot by 232%, and increased shoot biomass by 52%, but no effect was observed on total root system biomass. These results indicate that maize has a remarkable ability to maintain the total root system sink strength for carbon that leads to inherent reallocation of mass and length to embryonic roots (ER), first whorl nodal roots, and lateral roots of all classes. NR excision had no effect on shoot biomass in hydroponics, possibly showing that the root system was a stronger sink than the shoot for the excess available carbon. The increased shoot biomass, N content, and ^15^N content under low N in mesocosms indicates that the inherent modifications to root system architecture including greater root length of embryonic and all class lateral roots due to NRE leads to greater N uptake in solid media with positive effects on plant productivity. For the first time to our knowledge, a manipulative empirical study has confirmed that reduced nodal root number enhances N acquisition in maize under N replete conditions (York et al., 2013; Saengwilai et al., 2014) and has provided a mechanism through increased allocation to more efficient root classes.

Previous simulations using *SimRoot* indicated reduced NR number was a promising target for maize breeding (York et al., 2013; York, 2014), and these simulations were partially confirmed in greenhouse and field studies relying on genotypic variation within and among recombinant inbred line (RIL) populations (Saengwilai et al., 2014). Many NR-related quantitative trait loci (QTL) in maize have been reported for dry weight, length of NR (Burton et al., 2014), and approximately half of the total genetic variation in NR number was derived from QTLs for flowering time (Zhang et al., 2018). Though flowering time is related to NR number variation, the variation in NR number that *is not* due to flowering time will be more important for breeding. However, the NR-associated QTL on chromosomes have variation among maize populations, and no genes underlying the QTLs have been identified (Bray and Topp, 2018). Due to the fact maize RSA is controlled by many genes with generally small effects (Voss-Fels et al., 2018), genes like *rt1* (Jenkins, 1930), *rtcs* (Hetz et al., 1996) and *rtcl* (Taramino et al., 2007) in mutants of maize not only regulate NR formation, but also affect the development of other root classes and plant vigor, which means controlling NR number genetically for physiological experiments is difficult. Using root excision to manipulate NR number of maize may produce severe changes in metabolic processes, hormone homeostasis, and gas fluxes (Bloom and Caldwell, 1988), possibly like the reduced assimilation rate observed in the hydroponics study. Early changes in hormone levels in the rooting zone induced by root excision (Druege et al., 2016) and possible hormonal regulation of specific transporters to facilitate nutrient acquisition (Ghanem et al., 2011), were hypothesized to partially lead to increased shoot production in all experiments. However, no effect on shoot mass was observed in hydroponics due to NRE, therefore the fundamental relation of reduced nodal rooting to shoot mass likely is not due to hormonal differences or increased carbon availability. In fact, nodal root excision eliminates the confounding effects of many phenotypic differences that exist among genotypes, and can increase understanding of adventitious root formation and function (such as nodal roots), as well as allocation of carbon to the most efficient root classes (Poorter et al., 2012; Steffens and Rasmussen, 2016). Nodal root number in maize has been now investigated using simulations (York, 2014), field phenotyping of populations (Trachsel et al., 2011), physiological studies of contrasting lines for nodal root number (Saengwilai et al., 2014), and now a manipulative physiological study, which makes this phene aggregate one of the most complete examples of using the functional phenomics pipeline as outlined in York (2019). Therefore, NR number manipulation in large mesocosms using NRE is a valuable tool for better understanding the complex relations among roots, shoots, and soil resource uptake.

Carbon partitioning is governed by the relative source and sink strengths of all organs (Poorter et al., 2012), and there are trade-offs in carbon allocation between leaf, stem and root functions (McCarthy and Enquist, 2007). In general, plants should allocate carbon to various organs such that the marginal potential gain of carbon is equal to the marginal carbon allocation (Bloom et al., 1985). Early 6^th^ whorl NR formation was observed at 67% NRE under high N in mesocosms, implying NR initiation is partially determined by carbon availability. The 67% NRE level significantly decreased root mass faction by 12% in hydroponics (Fig. 1E) and by 19% in mesocosms (Fig. 4E), which suggests that these architectural changes led to more efficient N uptake throughout the experiment, resulting in greater shoot growth in LN. In turn, greater shoot mass may allow greater carbon fixation and subsequent allocation to both roots and shoots, causing positive feedback for plant growth, as predicted by York (2014). In general, no differences in overall root respiration and root system mass were observed due to NR excision, indicating a remarkable capacity of maize to maintain total root system sink strength and metabolic burden. For plants supplied with low N in mesocosms, the removal of 67% of nodal roots increased shoot biomass by 52% as compared to no excision control (Supplemental Table S2), and led to significant increases of root length in the deepest soil layer and increase of lateral-to-axial root length ratio across all the rooting depths except 90-120 cm, which could enhance greater soil exploration of heterogeneous supplies of nitrate in soil (Hodge, 2004), and may explain the enhanced plant growth by reducing NR number of plants in mesocosms with LN.

Specific root length is a good indicator of the relative ability to explore soil by roots, largely driven by thinner roots, and is strongly associated with the resource absorption function (Freschet et al., 2015). Removal 67% of NR significantly increased specific root length of NR5 under high N and NR3 and NR4 under low N in hydroponics and overall specific root length regardless of N levels in mesocosms, facilitated greater lateral-to-axial root length ratio of NR2, NR3 and NR4 in hydroponics and NR4 in mesocosms under low N, showing compensatory growth of lateral roots following root excision (Rubio and Lynch, 2007), which indicates enhanced capacity for resource acquisition (Lynch, 2013). Interestingly, this reallocation of NR carbon to lateral roots mimics root plasticity of maize under low N which may reduce NR number while increasing lateral root length (Gaudin et al., 2011). Maize lines with greater lateral root length have been shown to be more efficient at nitrogen capture (Zhan and Lynch, 2015). The thinner roots and greater lateral root length may favor root decomposition and buildup of soil carbon (Zhang and Wang, 2015), indicating possible ecosystem services beyond maintaining yield at lower nitrogen input levels.

Specific root respiration (SSR) by length and SRR by mass of the whole root system under high N were 2.0 and 2.6 times of those for roots under low N, which supports that root respiration is positively correlated with plant N concentration (Reich, 2014), and may be related to the changes in root morphological and biochemical phenes (Saglio and Pradet, 1980; Roumet et al., 2016). The reduction in SSR under low N could suggest it is adaptive in infertile conditions as a ‘cheap root’ phene state. Surprisingly, the deepest roots had greater SRR (mass based) regardless of N levels, possibly due to more axial or lateral root tips of per unit of root mass in the deepest layer. Young roots or growing root tips are important in resource acquisition at a high cost of energy consumption, and respiration declines with the aging of the roots (Wells and Eissenstat, 2003). This study indicates reduced specific root respiration could be an adaptive response to infertile soil conditions, which could possibly be affected by a multitude of root anatomical phenes that were not measured here (Burton et al., 2013). Overall, the results indicate that compensatory growth by embryonic, first whole nodal, and lateral roots of all root classes allowed the plant to maintain total metabolic burden of the root system while increasing soil exploitation under low N, but that further improvements could possibly be made by targeting reduced specific root respiration while increasing allocation to root classes with greater specific root length (cheap root classes).

## CONCLUSION

These results support the hypotheses that decreasing NR number increases the elongation of remaining of primary, seminal, nodal roots and their laterals, and increases N acquisition under low N condition in physical substrate. While compensatory growth of laterals was observed in the hydroponics study after nodal root excision, this did not lead to greater N uptake and shoot biomass, which indicates that excision of nodal roots doesn’t directly affect shoot biomass production as might be expected due to hormonal changes and overall plant carbon budget. Rather, our results indicate that by reallocating root mass to less expensive embryonic, early emerging nodal, and lateral roots throughout the soil profile, but especially at depth, plants with fewer nodal roots were able to more effectively exploit soil resources which led to greater N uptake and shoot growth. This research also highlights the importance of the embryonic root system due to its persistence over the course of the experiment and having the greatest rooting at depth relative to other axial classes in maize near flowering. Therefore, the reduced nodal root number ideotype is confirmed as a potential target for breeding programs, and a mechanism through greater soil resource uptake by laterals and early-emerging axial roots was revealed.

## MATERIALS AND METHODS

### Growth Chamber Hydroponics Study

#### Experimental Design

A hydroponics nutrient solution experiment was conducted in two growth chambers, arranged as a randomized complete block design replicated five times with a 2×2 factorial arrangement of treatments. The factors were two levels of N supply (high- and low-N conditions), and two levels of NR removal.

#### Growth Conditions

Black plastic pails (height 49.8 cm, top outside diameter 30.2 cm and bottom outside diameter 25.9 cm, with a volume of 28 L) with lids were placed in two Conviron E-15 growth chambers (Conviron, Winnipeg, Canada) with a day:night cycle of 14 h: 10 h, 28 °C: 22 °C, at a flux density at canopy level of c. 400 µmol m^−2^ s^−1^. Both chambers have same internal dimensions of 185 cm wide × 145 cm high × 79 cm deep. Seeds of maize inbred line B73 were obtained from Shawn Kaeppler, University of Wisconsin, Madison, WI, USA. Seeds were surface-sterilized in 0.05% NaOCl for 15 min and rinsed three times using DI water, then pre-germinated in medium size (0.3-0.5 mm) premium sand placed in darkness at 28 °C for 3 days, and transferred to growth chamber for 2 days. After that, uniformly germinated seedlings were washed out of the sand, wrapped around the junction between mesocotyl and coleoptiles with L800-D Identi-Plugs foam (Jaece Industries, NY), plugged in 5 cm Gro Pro net pot (Gro Pro, WA), and transplanted to a hole with diameter of 5 cm drilled into the lid with a hole saw. Each pail received one seedling, and the roots were directly submerged in nutrient solution. The nutrient solution for the high N treatment level was composed of (in *µ*M) 190 KH_2_PO_4_, 2260 KNO_3_, 750 CaCl_2_, 380 MgSO_4_, 17.29 H_3_BO_3_, 2.63 ZnSO_4_·7H_2_O, 3.38 MnCl_2_·4H_2_O, 0.12 CuSO_4_·5H_2_O, 0.04 (NH_4_)_6_Mo_7_O_24_·4H_2_O, and 300 Fe(III)-EDTA (C_10_H_12_N_2_N_a_F_e_O_8_). For the low N treatment level, KNO_3_ was reduced to 280 µM, and the differences between potassium supplies was balanced with KCl. The nutrient solution was continuously aerated with an air pump attached to airstones, and solution pH was maintained between 5.9 and 6.1 by additions of KOH or HCl throughout the experiment.

The nodal root (NR) number manipulation treatment levels included: excising two-third of all emerged nodal roots (67% NRE), and no nodal root excision control (0% NRE). NR removal was started 2 days after transplanting, plants were observed every other day due to the continuous emergence of nodal root, and roots were excised as needed to maintain the 67% target. Foam was gently squeezed to make counting of nodal roots emergence for both root excision and no root excision control treatments, and nodal roots were then excised close to its corresponded internode using scalpel for root excision treatment only. The targeted 67% NRE target was based on the calculation made from NR numbers of plants with excision.

##### Sample collection and measurements

One day before harvest, leaf gas exchange of the second youngest fully expanded leaf was measured in the growth chamber with a LI-6800 Portable Photosynthesis System equipped with the Multiphase Flash Fluorometer (LI-COR Inc., Lincoln, NE, USA). Leaf chamber conditions were as follows: fan speed of 10000 rpm, flow rate of 600 µmol mol^−1^, overpressure of 0.2 kPa, vapor pressure deficit at leaf of 1.2 kPa, leaf temperature of 25 °C, saturating irradiance of 1200 µmol m-^2^ s-^1^, and reference CO_2_ at 400 µmol mol-^1^. CO_2_ exchange was logged manually using stability criteria of both ΔH2O and ΔCO_2_ stdev limit of 0.1 over a period of 10 s.

Plants were harvested 24 days after transplanting when high N plants were at the 9-leaf stage and low N plants were at the 7-leaf stage. Shoots were dried at 60 °C for 3 days prior to dry weight determination. After a plant was removed from the nutrient solution, roots were immediately separated to the embryonic root system (including primary, seminal roots and their laterals) and different whorls of nodal roots, and each class was further divided into 30-cm sections along the axial roots of that class for an indication of rooting at various depths. All roots for a class and depth sample were blotted using tissue paper to remove excess water, then placed in a 19 ml custom chamber connected to the LI-8100 Automated Soil CO_2_ Flux System. The observation duration was 90 s, and the dead band was set at 20 s. All the respiration measurements were performed in an air-conditioned room with temperature maintained at 24 °C. After root respiration measurements, roots from each section were scanned as described below. Following scanning, the roots were dried at 60 °C for 3 days and weighed.

### Greenhouse Mesocosm Study

#### Experimental Design

The mesocosm experiment was conducted in a greenhouse of Noble Research Institute from July 30 to September 10, 2017, and arranged as a randomized complete block design replicated four times with a 2×3 factorial arrangement of treatments. The factors were two levels of N supply (high- and low-N conditions), and three levels of NR excision.

#### Growth Conditions

The mesocosm consisted of polyvinyl chloride (PVC) pipe (Charlotte, NC, USA) with a diameter of 15.24 cm and height of 152.4 cm, and flat bottom PVC cap (IPS Corporation, TN, USA). Each mesocosm was lined with seamless 6 mil heavy duty poly tubing (Uline, WI, USA) to facilitate root sampling at harvest. The growth medium consisted of mixture (volume based) of 50% medium size (0.3-0.5 mm) premium sand (Quikrete, GA, USA), 40% premium grade vermiculite (Sungro, MA, USA), and 10% perlite (Ambient Minerals, AR, USA). Each mesocosm was filled with 28 L of the mixture to ensure the same bulk density of the medium. The maize inbred line B73 seeds were obtained from Shawn Kaeppler, University of Wisconsin, Madison, WI, USA. Seeds were surface-sterilized in 0.05% NaOCl for 15 min and rinsed three times using DI water, and were then pre-germinated in rolled germination paper (Anchor Paper, MN, USA) soaked in 0.5 mM CaSO_4_ and placed in darkness at 28 °C in a germination chamber for 2.5 days. Two days before planting, each mesocosm received 6 L of nutrient solution. Each mesocosm received one seedling with the seed at a depth of 5 cm. Each plant was watered with 200 mL nutrient solution every other day. The nutrient solution for the high N treatment was composed of (in *µ*M) 500 KH_2_PO_4_, 6000 KNO_3_, 2000 CaCl_2_, 1000 MgSO_4_, 46 H_3_BO_3_, 7 ZnSO_4_·7H_2_O, 9 MnCl_2_·4H_2_O, 0.32 CuSO_4_·5H_2_O, 0.11 (NH_4_)_6_Mo_7_O_24_·4H_2_O, and 77 Fe(III)-EDTA (C_10_H_12_N_2_N_a_F_e_O_8_). For the low N treatment, KNO_3_ was reduced to 600 µM, and the differences between potassium supply was balanced with KCl. KOH was used to adjust nutrient solutions to pH 6.0. Three NR number manipulation treatments consisted of removing approximately one-third of all emerged nodal roots (33%), removing two-third of all emerged nodal roots (67%), and no root excision control (0%). NR excision was started 6 days after transplanting and continued every other day till the end of the experiment due to the continuous emergence of NR.

The growth media permitted root excision without damaging remaining roots. The approach of NR excision was to brush solid media away from the base of the stem where nodal roots emerged, use a scalpel to excise the necessary roots near their base, cover the stem again with the media, and give 200 ml nutrient solution to each plant in order to allow the media to settle. The targeted NRE levels of 33% or 67% were based on the calculations made from NR numbers of the corresponding plants with no excision. The no root excision control (0% NRE) received the same procedure for counting NR number over time, but none were excised.

#### Sample collection and measurements

The ability of roots to acquire N from deep soil layers was quantified using deep injection of ^15^NO_3_- (98% atom) 1 day before harvest. One hole was drilled at 143 cm depth in each mesocosm, and 5 ml of Ca^15^NO_3_ solution (0.46 mg ^15^N/mL) was injected into the hole of each mesocosm using a syringe. The plants were harvested at 42 d after planting. At harvest, the two newest fully expanded leaves were cut from the plant, the shoot was cut at the stem base, and all samples were dried at 60 °C for 3 days for biomass determination. The shoot samples were ground using Labman Automation Robot (Labman Automation, Stokesley, North Yorkshire, UK). The percents of total N and ^15^N in plant tissue were analyzed using a PDZ Europa ANCA-GSL elemental analyzer interfaced to a PDZ Europa 20-20 isotope ratio mass spectrometer (Sercon Ltd., Cheshire, UK) at the Stable Isotope Facility of the University of California at Davis (http://stableisotopefacility.ucdavis.edu/).

At harvest, a polyethylene liner in each mesocosm was carefully pulled out and the column of media and roots placed on a root washing station. Starting from the bottom, the liner was cut open and media were carefully washed away from roots using a low pressure water hose. After washing, roots were separated into embryonic roots (including primary, seminal roots, and their laterals) and different whorls of NR classes, and the roots of each class were divided into 30-cm sections along the axial roots for quantifying root distribution over depths for each class individually.

### Root Scanning and Image Analysis

For both the hydroponics and mesocosm experiments, roots from each class and depth sample were spread in a 5-mm layer of water in transparent plexiglass trays and imaged with a flatbed scanner equipped with a transparency unit (Epson Expression 12000XL, Epson America) at a resolution of 600 dpi. Images were analyzed using WinRhizo software (WinRhizo Pro, Regent Instruments, Quebec, CA) individually so that diameter thresholds could be adjusted for each image to distinguish axial roots from lateral roots, which would not be possible using batch mode. Root mass fraction was calculated as root dry weight proportion of total plant dry weight. Specific root length was calculated by dividing root length by the corresponding dry weight. The oven-dried root mass and root length quantified using WinRhizo were used to calculate the specific root respiration by root dry mass (SRR by mass; nmol g^−1^ s^−1^) and the specific root respiration by root length (SRR by mass; nmol cm^−1^ s^−1^), respectively. Individual axial root length was calculated by dividing total axial root length of each root class by the corresponding number of axial roots.

### Statistical Analysis

Statistical analyses were conducted using R version 3.5.1 (R Core Team, 2018) through RStudio version 1.1.45 (RStudio, 2016). Prior to data analysis, all the data were checked for normality with Shapiro-Wilk test and log-transformed when necessary. Regardless of data transformation, results presented in tables and figures report non-transformed values. Analysis of variance (ANOVA) was performed across nitrogen (N) levels and nodal root excision (NRE) levels for comparisons between N levels, NRE levels, and their interaction based on the phenes measured at the whole plant level (not separated by root class or depth). N levels, NRE levels and their interactions were analyzed as fixed effects in a mixed effects model with block as random effects. Root distribution data across root classes and rooting depths were analyzed with one-way ANOVA under each N level separately, and with pairwise comparisons of NRE levels under either same root classes or same rooting depths, because the focus was on NRE levels and otherwise the models were difficult to interpret and report. Tukey’s test at *α* = 0.05 was used for multiple comparisons among different treatments. The R packages ‘emmeans’ (Lenth, 2018), ‘lme4’ (Bates et al., 2014), ‘multcompView’ (Graves et al., 2012), were used for statistical analyses and the R package ‘ggplot2’ (Wickham, 2016) for data visualization.

### Supplemental Data

The following supplemental materials are available.

## ACKNOWLEDGMENTS

We thank Bryce Walker, Anand Seethepalli, Na Ding for assistance sampling and root scanning, and David Huhman and Bonnie Watson for their technical assistance for analyzing nitrogen content of samples. This research was supported by the Noble Research Institute, LLC and the Samuel Roberts Noble Foundation.

---

**Supplemental Figure S1.**
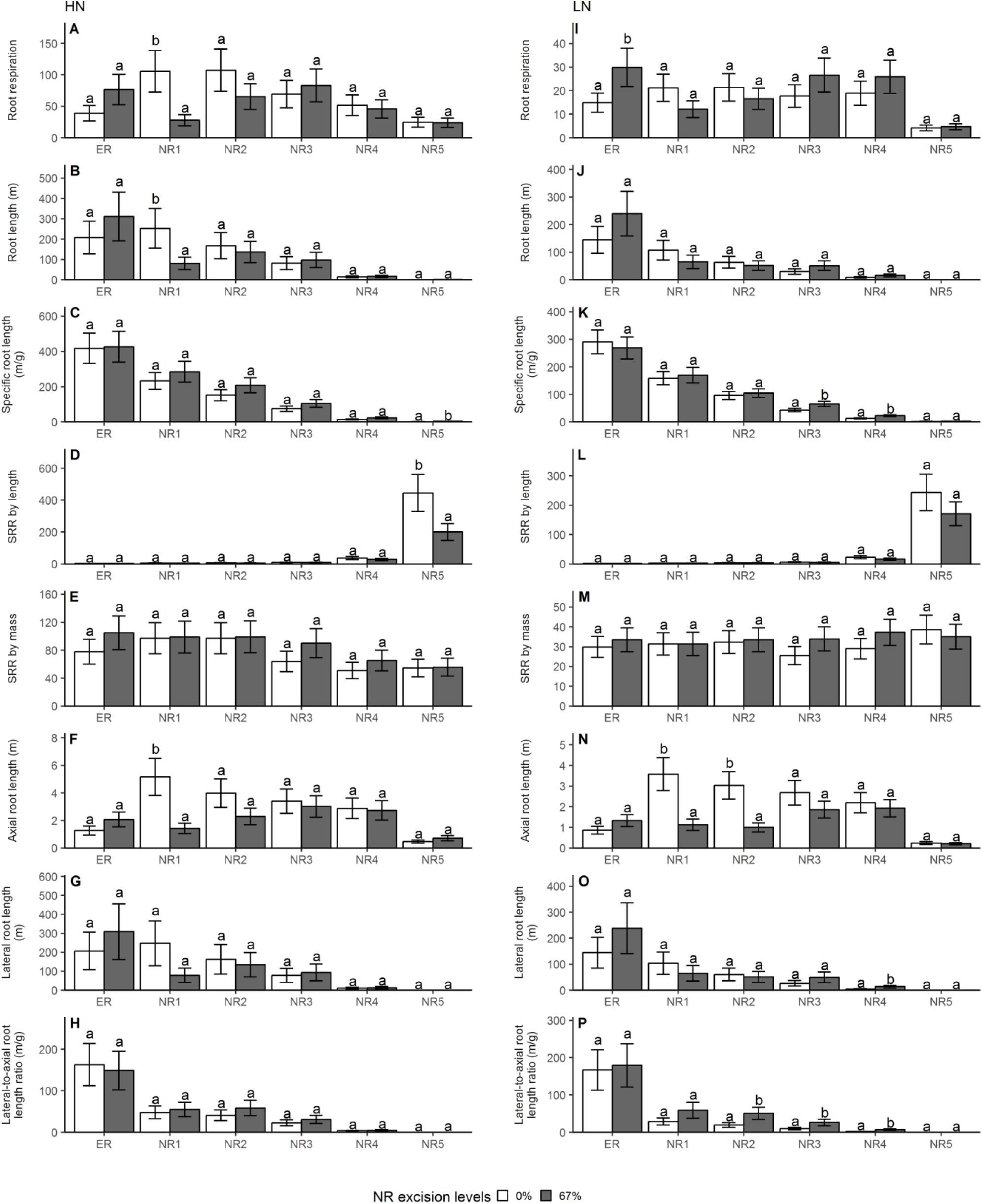
Effect of NR excision treatment on root phenes of different root classes under high N (A, B, C, D, E, F, G, H) and low N (I, J, K, L, M, N, O, P) level in growth chamber hydroponics study. Each column represents the mean of n=5 for each treatment. Error bars represent standard errors, and columns with the same letter under same root class were not significantly different at p≤0.05 according to Tukey’s test. ER: embryonic roots, NR1: 1^st^ whorl nodal roots, NR2: 2^nd^ whorl nodal roots, NR3: 3^rd^ whorl nodal roots, NR4: 4^th^ whorl nodal roots, NR5: 5^th^ whorl nodal roots, SRR: specific root respiration.

**Supplemental Figure S2.**
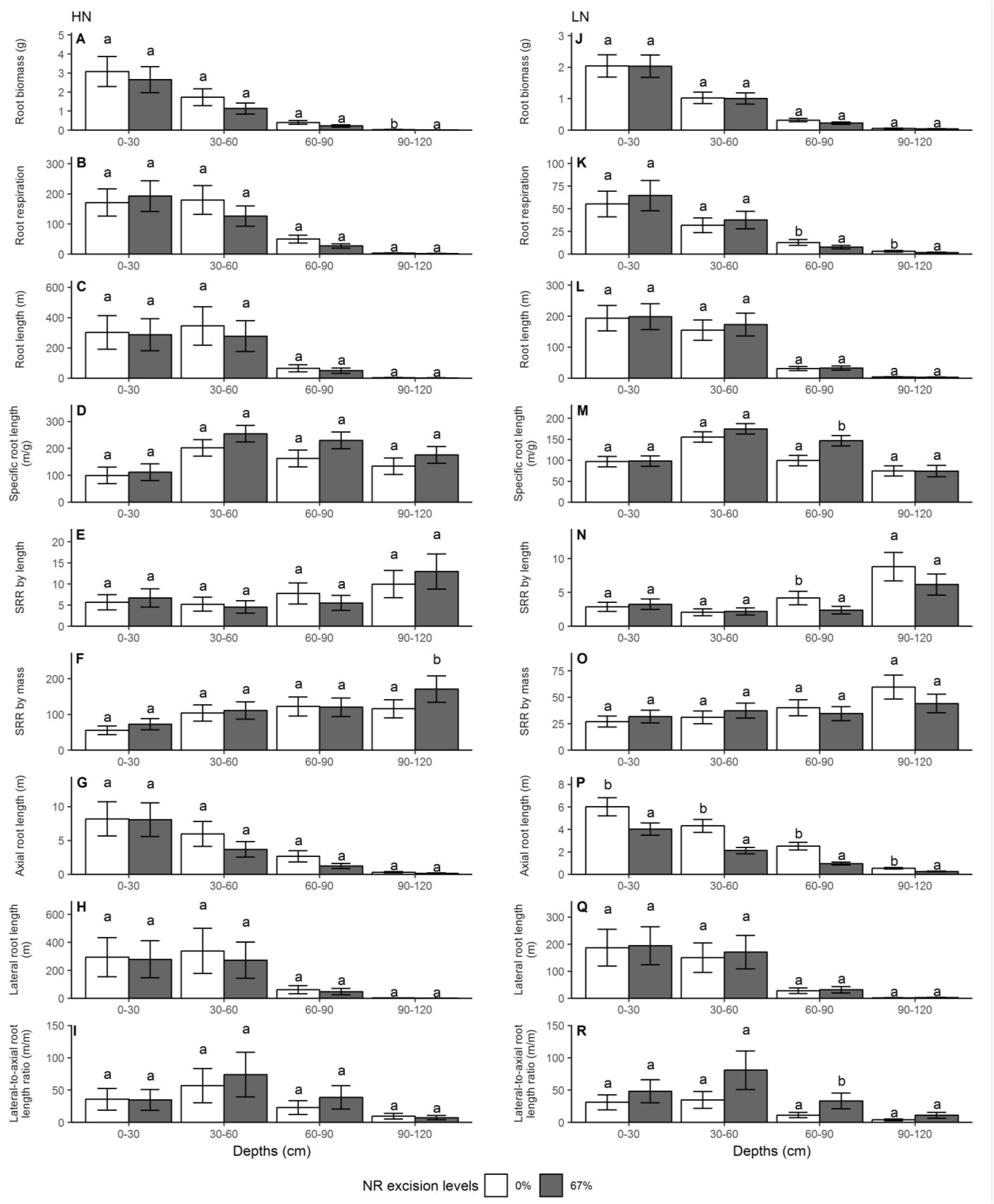
Effect of NR excision treatment on root phenes of different rooting depths under high N (A, B, C, D, E, F, G, H, I) and low N (J, K, L, M, N, O, P, Q, R) level in growth chamber hydroponics study. Each column represents the mean of n=5 for each treatment. Error bars represent standard errors, and columns with the same letter under same depth were not significantly different at p≤0.05 according to Tukey’s test. SRR: specific root respiration.

**Supplemental Figure S3.**
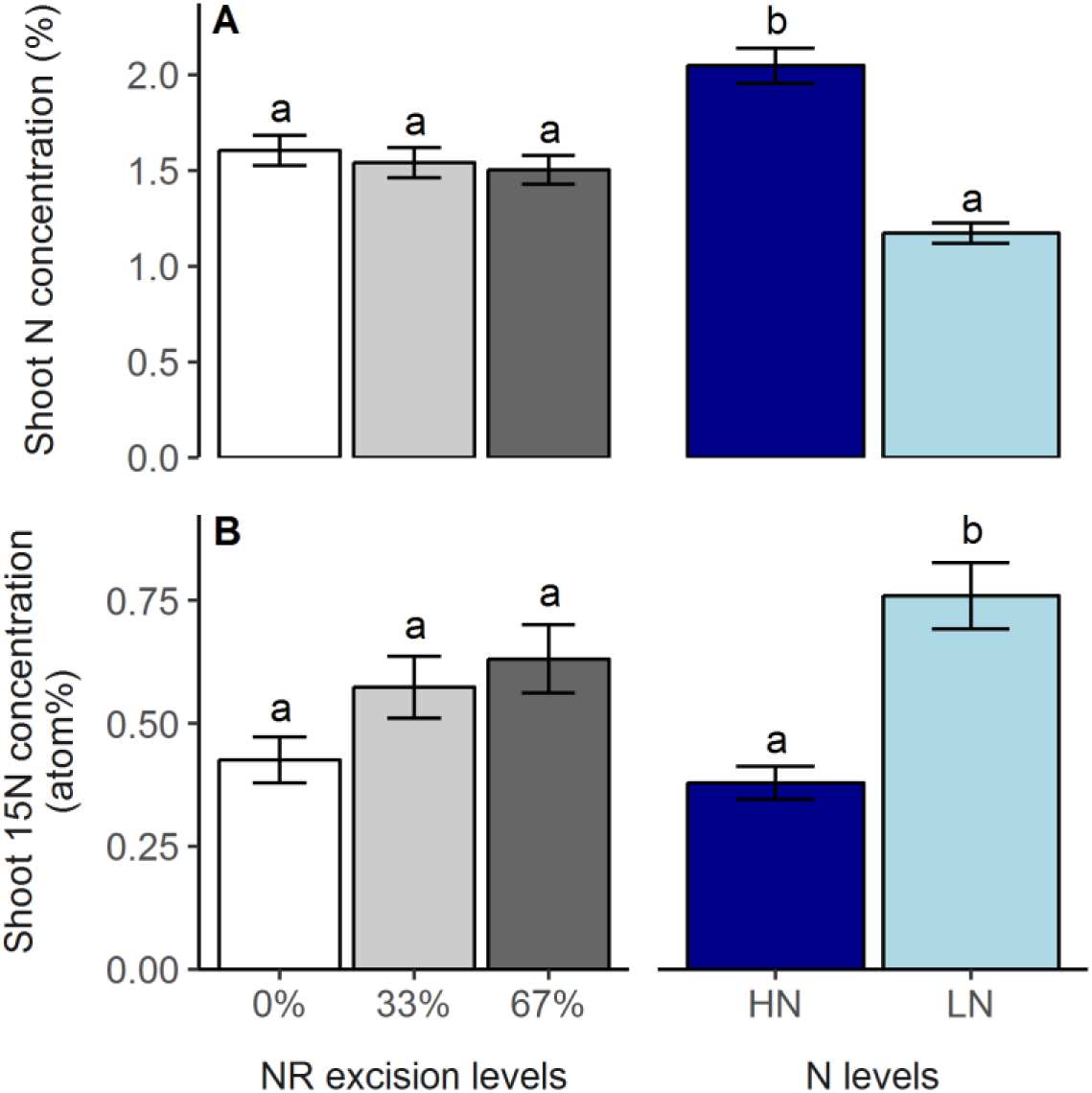
Main effects of N treatment and NR excision treatment on shoot N concentration (A) and shoot ^15^N concentration (B) for the greenhouse mesocosms. Each column represents the mean of n=4 for each treatment. Error bars represent standard errors, and columns with the same letter under same main effect were not significantly different at p≤0.05 according to Tukey’s test.

**Supplemental Figure S4.**
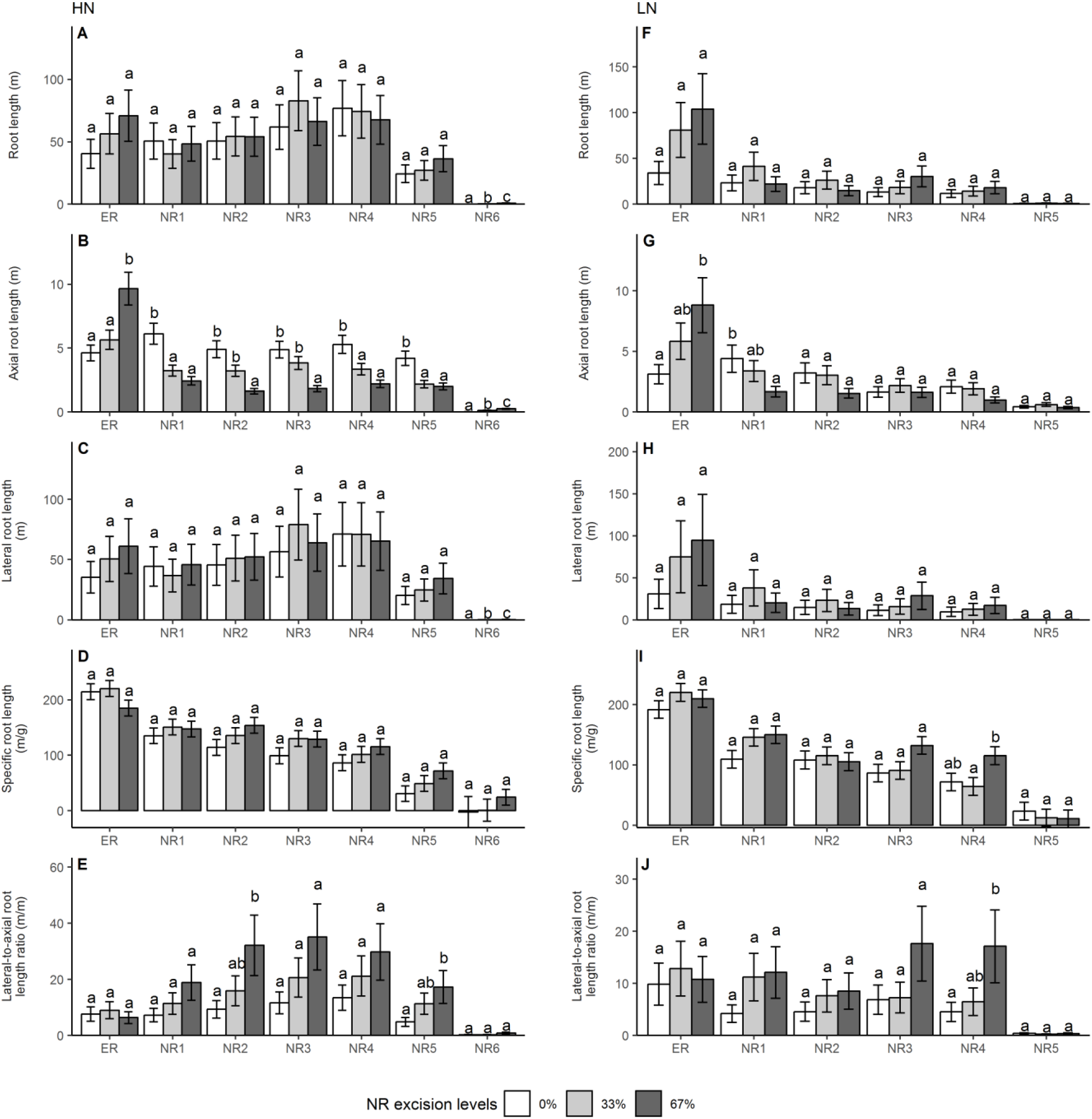
Effect of NR excision treatment on root phenes of different root classes under high N (A, B, C, D, E) and low N (F, G, H, I, J) level in in the greenhouse mesocosms. Each column represents the mean of n=4 for each treatment. Error bars represent standard errors, and columns with the same letter under same root class were not significantly different at p≤0.05 according to Tukey’s test. ER: embryonic roots, NR1: 1^st^ whorl nodal roots, NR2: 2^nd^ whorl nodal roots, NR3: 3^rd^ whorl nodal roots, NR4: 4^th^ whorl nodal roots, NR5: 5^th^ whorl nodal roots, NR6: 6^th^ whorl nodal roots.

**Supplemental Figure S5.**
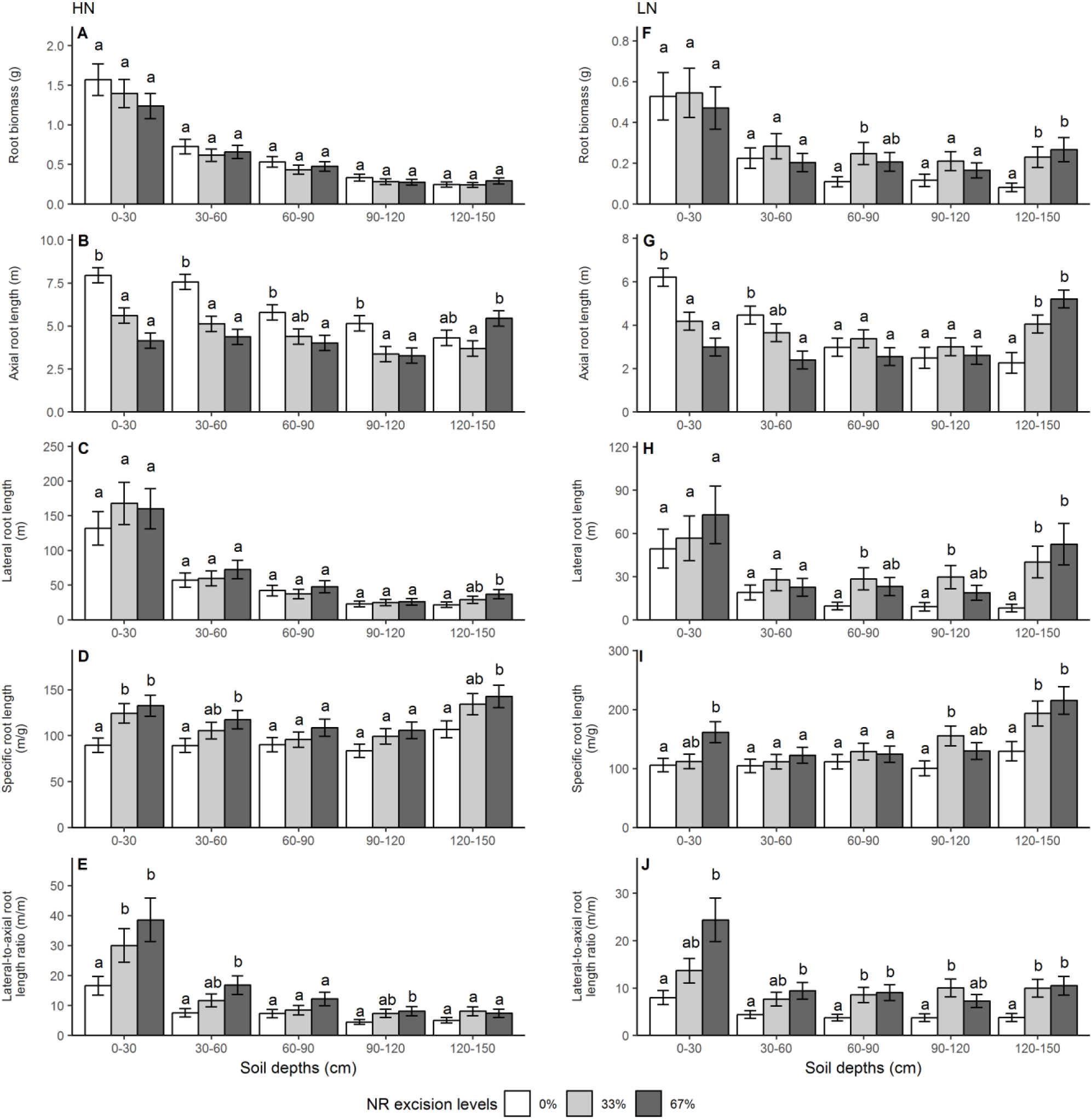
Effect of NR excision treatment on root phenes of different rooting depths under high N (A, B, C, D, E) and low N (F, G, H, I, J) level in greenhouse mesocosms study. Each column represents the mean of n=4 for each treatment. Error bars represent standard errors, and columns with the same letter under same depth were not significantly different at p≤0.05 according to Tukey’s test.

**Supplemental Table S1.** Summary of ANOVA (F values and significance lvels) of whole-plant shoot and root phenes as influenced by N treatment, NR excision treatment and their interactions for the two experiments.

| Phenes | Factors |  |  |
| --- | --- | --- | --- |
|  | N treatment (N) | NR excision treatment (NRE) | N*NRE |
| ***Growth chamber hydroponics*** |  |  |  |
| Nodal root number | 7.98 * | 211.65 *** | 0.01 ns |
| Shoot biomass | 38.35 *** | 0.09 ns | 0.27 ns |
| Root biomass | 7.61 * | 1.24 ns | 0.39 ns |
| Total biomass | 28.07 *** | 0.01 ns | 0.30 ns |
| Root mass fraction | 4.22 ns | 5.11 * | 0.07 ns |
| Assimilation rate | 782.17 *** | 10.87 ** | 0.03 ns |
| Transpiration rate | 185.84 *** | 0.16 ns | 0.00 ns |
| Stomatal conductance | 157.31 *** | 0.05 ns | 0.08 ns |
| Total root length | 62.74 *** | 0.44 ns | 0.03 ns |
| Specific root length | 11.21 ** | 4.71 * | 1.11 ns |
| Specific root respiration by length | 66.50 *** | 0.24 ns | 0.19 ns |
| Specific root respiration by mass | 120.55 *** | 2.37 ns | 0.03 ns |
| Axial root length | 25.69 *** | 17.13 ** | 1.67 ns |
| Lateral root length | 61.76 *** | 0.59 ns | 0.04 ns |
| Lateral-to-axial root length ratio | 1.08 ns | 19.62 *** | 3.41 ns |
| ***Greenhouse mesocosms*** |  |  |  |
| Nodal root number | 82.69 *** | 339.33 *** | 1.83 ns |
| Shoot biomass | 164.30*** | 4.08* | 2.21 ns |
| Root biomass | 66.02*** | 0.41 ns | 1.85 ns |
| Total biomass | 140.91*** | 2.80 ns | 2.11 ns |
| Root mass fraction | 29.36*** | 9.89** | 0.06 ns |
| Shoot N concentration | 156.97*** | 0.75 ns | 0.28 ns |
| Shoot <sup>15</sup>N concentration | 30.31 *** | 3.51 ns | 3.21 ns |
| <sup>15</sup>N in shoot | 43.12 *** | 6.13 ** | 5.44 * |
| N in shoot | 1087.62*** | 7.55** | 6.95** |
| Total root length | 32.65*** | 2.44 ns | 0.25 ns |
| Specific root length | 13.19** | 13.58*** | 0.75 ns |
| Axial root length | 42.90*** | 7.78** | 5.14* |
| Lateral root length | 30.80*** | 2.91 ns | 0.19 ns |
| Lateral-to-axial root length ratio | 11.37** | 12.62*** | 0.03 ns |

\*, \*\*, and \*\*\* significant at 0.05, 0.01 and 0.001 levels, respectively. ns = not significant at 0.05 level.

**Supplemental Table S2.** Mean ±SE of shoot biomass as influenced by NR excision treatment under low N conditions in the greenhouse mesocosms experiment.

| **N level** | **NR excision levels** | **Mean ± SE** | **Phenes** |
| --- | --- | --- | --- |
| LN | 0% | 3.76±1.18 | Shoot biomass (g) |
| LN | 33% | 5.58±0.35 | Shoot biomass (g) |
| LN | 67% | 5.72±0.79 | Shoot biomass (g) |

Author contributions
H.G performed all of the experiments, analyzed the data, and wrote the article with contributions from the corresponding author; L.M.Y conceived the hypotheses and supervised the design, experimentation, data analysis, and reporting.

